# Using supervised learning methods for gene selection in RNA-Seq case-control studies

**DOI:** 10.1101/282780

**Authors:** Stephane Wenric, Ruhollah Shemirani

## Abstract

Whole transcriptome studies typically yield large amounts of data, with expression values for all genes or transcripts of the genome. The search for genes of interest in a particular study setting can thus be a daunting task, usually relying on automated computational methods. Moreover, most biological questions imply that such a search should be performed in a multivariate setting, to take into account the inter-genes relationships.

Differential expression analysis commonly yields large lists of genes deemed significant, even after adjustment for multiple testing, making the subsequent study possibilities extensive.

Here, we explore the use of supervised learning methods to rank large ensembles of genes defined by their expression values measured with RNA-Seq in a typical 2 classes sample set. First, we use one of the variable importance measures generated by the random forests classification algorithm as a metric to rank genes. Second, we define the EPS (extreme pseudo-samples) pipeline, making use of VAEs (Variational Autoencoders) and regressors to extract a ranking of genes while leveraging the feature space of both virtual and comparable samples.

We show that, on 12 cancer RNA-Seq data sets ranging from 323 to 1210 samples, using either a random forests based gene selection method or the EPS pipeline outperforms differential expression analysis for 9 and 8 out of the 12 datasets respectively, in terms of identifying subsets of genes associated with survival.

These results demonstrate the potential of supervised learning-based gene selection methods in RNA-Seq studies and highlight the need to use such multivariate gene selection methods alongside the widely used differential expression analysis.

## 1 Introduction

Transcriptomics studies making use of RNA-Seq usually produce large amounts of data, namely one expression value for each gene or transcript of each sample assessed [Wang2009, Mortazavi2008].

Searching for genes of interest or prioritizing genes in the context of case-control studies related to diseases or other experimental conditions constitutes an important task ascribed to RNA-Seq experiments [Trapnell2009, Garber2011, Love2014, Wenric2017].

Current methods often make use of differential expression analysis, to select genes of interest and assign them a p-value related to a statistical test assessing changes in expression between different conditions.

Most commonly used software packages performing differential expression analysis make use of the negative binomial distribution to model read counts for each gene. This distribution, which is an extension of the Poisson distribution, has two parameters: the mean and the dispersion, which allows modeling of more general mean–variance relationships than Poisson. The dispersion parameter allows to take into account the biological variability arising in RNA-Seq data [Love2014, Huang2015].

However, even though software packages like DESeq2 model relationships between genes by assuming that genes of similar average expression have a similar dispersion, the statistical test conducted to assess significance is a univariate test performed independently for each gene. Albeit providing particularly useful and usually accurate information regarding disruptions of gene expression between conditions, these methods thus do not take into account the potential correlation and concordant or discordant effect between groups of genes. However, such gene-gene interactions are present in most tissues and conditions and they are known to play key roles in said conditions, with groups of genes which might have a significant effect as a group but not when each gene is considered independently [Kanehisa2000, Joshitope2005, Phillips2008, Vidal2011].

Here, we explore the use of multivariate classifiers to rank genes in a case-control RNA-Seq experiment. Namely, we’re using the permutation importance of the random forests classifier to rank genes, and a newly developed method (EPS) making use of Variational Autoencoders.

Machine learning methods are progressively being applied to problems arising in genomics related fields and the idea of using importance measures generated by the random forests algorithm to extract a ranking of features has already been explored with several different data sets, although, to our knowledge, this has never been done with RNA-Seq data sets [Schrider2018, Freres2016, Yao2015, Duro2012, Anaissi2013].

Aside from random forests, we also introduce a technique called Extreme Pseudo-Sampling (EPS) allowing to create case and control pseudo-samples lying on the two extremes of the sample space. This method uses Variational Autoencoders (VAE) [Kingma2013] to create new pseudo-samples that are not present in the original datasets but closely imitate their statistical properties, in that they share the properties of independent and identically distributed samples from the same distribution as the real data.

The idea of using autoencoders to classify and examine genomics datasets is not new [Tan2015]. However, VAEs differ from other autoencoders in that they can create a meaningful latent representation space where one can choose a new vector in the latent space and create a valid, previously unseen sample in real space that closely follows the real samples (the aforementioned pseudo-samples).

Additionally, although autoencoders have been used as an auxiliary tool in the classification of existing datasets, no attempt has been made to extract the knowledge learnt by the autoencoders in this process to trace the analysis and results back to the actual gene expression values and their relationships. Here, we suggest a way to make use of that information [Tan2015].

## 2 Materials and Methods

### 2.1 Data sets

Several data sets from the TCGA database have been selected to validate both methods [Weinstein2013].

Only the data sets containing 30 healthy samples (denoted as “Solid Tissue Normal” in the TCGA database) or more have been selected. All read counts produced by HTSeq as well as the clinical data have been downloaded with the TCGABiolinks R/Bioconductor package [Colaprico2016].

The data sets selected are summarized in Table 1.

**Table 1.** TCGA data sets used in this study.

| Name | Cancer type | N (tumors) | n (healthy) | Median age | Age range |
| --- | --- | --- | --- | --- | --- |
| TCGA-BRCA | Breast invasive carcinoma | 1097 | 113 | 59.07 | 26-90 |
| TCGA-LUAD | Lung adenocarcinoma | 582 | 59 | 66.88 | 33-88 |
| TCGA-UCEC | Uterine Corpus endometrial carcinoma | 559 | 35 | 64.24 | 31-90 |
| TCGA-KIRC | Kidney renal clear cell carcinoma | 535 | 72 | 61.16 | 26-90 |
| TCGA-HNSC | Head and neck squamous cell carcinoma | 528 | 44 | 61.14 | 20-90 |
| TCGA-THCA | Thyroid carcinoma | 507 | 58 | 46.92 | 15-89 |
| TCGA-LUSC | Lung squamous cell carcinoma | 504 | 49 | 68.66 | 39-90 |
| TCGA-PRAD | Prostate adenocarcinoma | 498 | 52 | 61.99 | 42-78 |
| TCGA-COAD | Colon adenocarcinoma | 460 | 41 | 68.88 | 31-90 |
| TCGA-STAD | Stomach adenocarcinoma | 443 | 32 | 67.56 | 30-90 |
| TCGA-LIHC | Liver hepatocellular carcinoma | 377 | 50 | 61.53 | 16-88 |
| TCGA-KIRP | Kidney renal papillary cell carcinoma | 291 | 32 | 62.03 | 28-88 |

### 2.2 Methodology

For each data set, the methodology illustrated in Fig. 1 has been applied:

- All samples are normalized with the DESeq2 software package [Love2014].
- The samples are split into a training set and a validation set. The training set contains all the healthy samples of the original data set (*n*) and the same number of tumor samples as healthy samples (*n*). The validation set contains the remaining tumor samples (*N - n*).
- Differential expression analysis is performed on the training set with the DESeq2 software package, using default parameters and options. A ranking of genes, based on their adjusted p-value relative to the differential expression test, is obtained.
- A random forests classifier is built on the training set with the ranger R package, using 100000 trees and a value for the *m*_*try*_ parameter of 236 (equal to the square root of the total number of features) [Wright2015]. A ranking of genes based on their permutation importance values is obtained (the permutation importance is computed by randomly permuting the values of the feature of interest and measuring the resulting increase in error).
- The Extreme Pseudo-Sampling method (see 2.3) is applied on the training set(s) to extract a ranking of genes.
- Let *RF* denote the random forests based gene ranking, *DE* the differential expression based gene ranking and *EPS* the extreme pseudo-samples based gene ranking. *RF*_*i*_ denotes the *i*-th gene of the random forests based gene ranking. Similarly, *DE*_*i*_ denotes the *i*-th gene of the differential expression based gene ranking and *EPS*_*i*_ denotes the *i*-th gene of the extreme pseudo-samples based gene ranking.
- For both rankings, 20 gene signatures are generated, including an incremental number of genes. Let *sigRF*_*i*_ denote the *i*-th gene signature based on the random forests ranking, *sigDE*_*i*_ denote the *i*-th gene signature based on the differential expression ranking and *sigEPS*_*i*_ the *i*-th gene signature based on the extreme pseudo-samples ranking. The signatures are formally defined as:
  - *sigRF*_*i*_ = {*RF*_1_,…,*RF*_*i*_}, for *I* = 1…20
  - *sigDE*’ = {*DE*_1_…*DE*_*i*_}, for *i*= 1…20
  - *sigEPS*_*i*_ = {*EPS*_1_…*EPS*_*i*_}, for *i*= 1…20
- For each signature,
  - A Cox proportional hazard model was built using all genes of the signature
  - The samples of the validation set were split into two groups (higher and lower survival), based on the median of the Cox proportional hazard model.
  - A log-rank test was performed to compare the survival of the two groups.
- For *i* = {1, …, 20}, the p-value of the log-rank tests obtained with *sigDE*_*i*_, *sigRF*_*i*_, *sigEPS*_*i*_ are compared.

For each data set, correlation coefficients have been computed between the expression values of the 50% most expressed genes; a hierarchical clustering of the 50% most expressed genes was performed, to assess if multicollinearity played a role in the performance of the RF based method (multicollinearity denotes the presence of non-independent features such that the relationship between each of these features and the model output is influenced by the relationships between the non-independent features). A hierarchical clustering of all samples was also performed, with the 50% most expressed genes. Enrichment analysis was performed on gene lists from both methods.

The correlation coefficient between each top-ranked gene from both list and the 50% most expressed genes has been computed for each data set.

Globally, the correlation between the overall survival at 5 years of all cancer types, and the performance of the presented methods was computed.

### 2.3 Extreme Pseudo-Sampling

It is worth noting that, in most data sets considered in this study, the samples from both classes reside in a high dimensional space and are tightly coordinated together, such that a linear classifier cannot separate them at all. The low count of normal samples compared to the total sum of samples also contributes to the failure of linear classifiers; which tend to receive bias from such unbalance of class membership statistics.

We decided to use a dimensionality reduction technique in order to both address the *curse of dimensionality* and find a representation in which these samples lay in a linearly-separable subspace.

Autoencoders have shown to be able to create such latent representations better than their linear counterparts such as PCA [Tan2015, Danaee2016]. However, such representations do not provide us with useful, actionable knowledge about genes due mainly to their non-linear activation functions.

Moreover, Normal Autoencoders are not generative, i.e. while it is possible to come up with useful latent representations for classification purposes, one cannot generate new samples similar to the real samples by slightly modifying their latent representation values and feeding the result into the decoder network.

A new type of Autoencoder, called the Variational Autoencoder, however, can succeed in this task [Kingma2013]. VAEs are fundamentally different from other AEs in that they are generative models:

Each point *x* in real space will be associated with distribution *P(z*|*x)*. For the purpose of this methodology, we assumed this distribution to be normal. Getting latent representation *z*_*1*_ from sample *x*_*1*_, thus, would be equal to drawing a sample from distribution *𝒩(µ*_*1*_, *σ*_*1*_), where *µ*_*1*_, *σ*_*1*_ are learned from the training data.

The training VAE comprises 9 layers, having 30000, 15000, 10000, 2000, 500, 2000, 10000, 15000, 30000 perceptrons respectively. The training process of these layers requires fine-tuning approximately 5 billion parameters. Given that the performance of this fine-tuning process increases with the number of samples, in addition to the training set extracted from the studied TCGA dataset, a random selection of samples from the 11 other training sets is used in the VAE training process.

After the training step, each dataset *D*_*c*_ is transformed to its latent representation *L*_*c*_. Said latent representation allows to linearly separate the normal samples from cancerous ones with almost 100% accuracy for both testing and training datasets. Considering the linear separator, let us denote the furthest populated areas on both sides of the separator, called *N*_*c*_ for the normal side of the linear separator and *C*_*c*_ for the cancerous side. If we consider a point *z*_*n*_ in one of these areas, we know it has been randomly drawn from distribution *𝒩(µ*_*n*_, *σ*_*n*_).

While selecting *z*_*n*_ is a random process, once a *z*_*n*_ has been drawn from any of the distributions, reconstructing 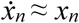 from *z*_*n*_ is a deterministic process done by the decoder. However, every point in the close proximity of *z*_*n*_ can be drawn from the same distribution. Due to the deterministic features of the decoder, each of these points would end up generating a different 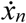. Although different, every possible 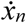 should resemble the original *x*_*n*_ closely and should also follow the general statistical characteristics of all *x*’s in the dataset.

We then drew 400 random points in areas *N*_*c*_ *and C*_*c*_ of the latent space *L*_*c*_, on both sides of the linear separator and generated new “virtual” or “pseudo” samples of both cancerous and normal classes, a process that we call Extreme Pseudo Sampling (EPS). The amount of random points drawn (400) was chosen using cross validation on the training data. It was the smallest number of samples that ended up in a successful regression process.

While real samples cannot be divided using a linear separator and suffer from unbalance of class member counts; we were able to generate new pseudo samples that can be divided linearly in real space due to their exaggerated cancerous/normal features. These samples also are of equal count. The later trait enables the dividing regression lines to be less biased towards a specific class. Thus, said regression lines maintain the same distance from both classes.

Finally, since all sample features have been normalized in the process, weight coefficients in the line formula can be translated into importance factors for classifying extreme pseudo samples. The larger a coefficient, the more important its related feature is in determining class membership. Thereby, we are able to extract an importance ranking for all genes, in each data set.

The R and Python scripts used to perform the aforementioned analyses are available online: https://github.com/stephwen/ML_RNA-Seq & https://github.com/roohy/Extreme-Pseudo-Sampler

## 3 Results

For each data set, 60 log-rank tests have been performed on the validation set, using gene signatures *sigDE*_*i*_, *sigRF*_*i*_, and *sigEPS*_*i*_ with *i = {1, 2*, …, *20}* which contain from 1 to 20 genes out of the gene ranking derived from differential expression analysis, the gene ranking derived from the random forests classifier, and the gene ranking derived from the Extreme Pseudo-Sampling method respectively. The p-values of these tests have been compared two by two.

Table 2 summarizes the results and shows the number of gene signatures where the random forests based gene ranking outperforms the differential expression based gene ranking and where the Extreme-Pseudo Sampling method outperforms the differential expression based gene ranking.

**Table 2.** The random forests column denotes the number of random forests based signatures having a lower log-rank p-value than their corresponding differential expression based signatures. The extreme pseudo-samples column denotes the number of extreme pseudo-samples based signatures having a lower log-rank p-value than their corresponding differential expression based signatures. The 3 colors (green, yellow, red) refer to cases where the proposed methods have a higher number, the same number, and a lower number of best-performing gene signatures than DESeq2, respectively.

| Name | Cancer type | Random forests | Extreme pseudo-samples |
| --- | --- | --- | --- |
| TCGA-BRCA | Breast invasive carcinoma | 5 | 19 |
| TCGA-LUAD | Lung adenocarcinoma | 14 | 14 |
| TCGA-UCEC | Uterine Corpus endometrial carcinoma | 16 | 9 |
| TCGA-KIRC | Kidney renal clear cell carcinoma | 13 | 10 |
| TCGA-HNSC | Head and neck squamous cell carcinoma | 14 | 15 |
| TCGA-THCA | Thyroid carcinoma | 15 | 15 |
| TCGA-LUSC | Lung squamous cell carcinoma | 5 | 0 |
| TCGA-PRAD | Prostate adenocarcinoma | 12 | 19 |
| TCGA-COAD | Colon adenocarcinoma | 11 | 18 |
| TCGA-STAD | Stomach adenocarcinoma | 13 | 19 |
| TCGA-LIHC | Liver hepatocellular carcinoma | 19 | 8 |
| TCGA-KIRP | Kidney renal papillary cell carcinoma | 10 | 19 |

For <u>9</u> out of the 12 data sets analyzed (lung adenocarcinoma, uterine corpus endometrial carcinoma, kidney renal clear cell carcinoma, head and neck squamous cell carcinoma, thyroid carcinoma, prostate adenocarcinoma, colon adenocarcinoma, stomach adenocarcinoma, liver hepatocellular carcinoma), the random forests based gene ranking outperforms the differential expression based gene ranking in terms of identifying subsets of genes associated with survival._For 8 out of the 12 datasets (breast invasive carcinoma, lung adenocarcinoma, head and neck squamous cell carcinoma, thyroid carcinoma, prostate adenocarcinoma, colon adenocarcinoma, stomach adenocarcinoma, kidney renal papillary cell carcinoma), the extreme pseudo-samples based gene ranking outperforms the differential expression based gene ranking. For one data set (kidney renal papillary cell carcinoma), both the DESEq2 and the random forests based gene rankings share the same number of best performing signatures. For one data set (kidney renal clear cell carcinoma), both the DESEq2 and the extreme pseudo-samples based gene rankings share the same number of best performing signatures. For 2 out of the 12 data sets (breast invasive carcinoma, lung squamous cell carcinoma), the differential expression based gene ranking outperforms the random forests based gene ranking. For 3 out of the 12 data sets (uterine corpus endometrial carcinoma, lung squamous cell carcinoma, liver hepatocellular carcinoma), the differential expression based gene ranking outperforms the extreme pseudo-samples based gene ranking.

Figure 2 shows the log-rank p-values for the 3 different methods (DESeq2, random forests, extreme pseudo-samples) and their respective gene signatures ranging from 1 to 20 genes, for the 4 largest data sets (TCGA-BRCA, TCGA-LUAD, TCGA-UCEC, TCGA-KIRC). Similar figures for the 8 other data sets are available as supplementary data. The log-rank p-values for the 20 gene signatures related to the 3 rankings for each dataset and the genome wide ranking of genes based on the permutation importance computed by the random forests classifier and on the extreme pseudo-samples method can be found in Supplementary Table 1 and Supplementary Table 2 respectively.

No significant difference in the average absolute correlation coefficient obtained between the 50% most expressed genes was found between the different cohorts whose DE based signatures performed better than the RF and EPS signatures and the cohorts whose RF or EPS based signatures performed better than the DE ones._No significant difference in terms of the number of clusters of samples obtained with a hierarchical clustering with the 50% most expressed genes when using a constant height cutoff value of *h = 2*\**10*^*6* was found between the different cohorts whose DE based signatures performed better than the RF and EPS signatures and the cohorts whose RF or EPS based signatures performed better than the DE ones._No significant difference in terms of the number of clusters of genes obtained with a hierarchical clustering with the 50% most expressed genes when using a constant height cutoff value of h = 10^5 was found either._No significant difference was found between the correlation between the top-ranked genes selected with both methods and the 50% most expressed genes. No correlation was found between the overall survival at 5 years of the different cancer types and the performance of either method (measured as the ratio of n/20 top-performing signatures). There is, however, a loose correlation (Pearson correlation coefficient: 0.627, p-value: 0.029) between the number of best-performing DE based signatures among the 20 signatures of each data set and the number of differentially expressed genes (adjusted p-value < 0.05) in each data set. Correlation coefficients and numbers of clusters are present, for all data sets, in Supplementary Table 3.

## 4 Discussion

Highlighting genes of interest has always been a part of transcriptomics studies and the advent of RNA sequencing technologies has but further emphasized this endeavor. Traditionally, genes of interest, in case-control studies where one had access to their expression values, were genes where said expression varied greatly from one class to the other. This definition has led to the development of numerous methods making use of diverse statistical models and tests, achieving impressive results in a lot of different use cases. However, these methods often implicitly neglected the importance of gene-gene relationships, by only looking at univariate changes.

Here, we propose a paradigm shift, by directing the search for genes of interest towards the use of machine learning methods originally conceived to predict the membership of a sample in a class, as these methods intrinsically model the inter-variable relationships (*i*.*e*. the previously overlooked gene-gene links).

An obvious kind of data sets which should theoretically benefit from this are cancers, as these pathologies are known to involve several genes in a multistep process, with different mechanisms implicating intricate relationships between said genes [Vogelstein2013, Yates2012].

By using 12 data sets containing samples of various cancers, we have shown that supervised classification algorithms could be used to extract a meaningful ranking of genes. Namely, the permutation importance (also known as Mean Decrease in Accuracy) generated by the random forests algorithm and the weights coefficients used in the extreme pseudo-samples provided a ranking of genes which outperformed classical methods in most data sets.

The permutation importance is not the only variable importance generated by the random forests classifier, as the Gini importance (or Mean Decrease in Impurity) is also available. However, using the Gini importance to classify the genes of these data sets yielded slightly worse results than the results obtained with the permutation importance. Using a combination of both variable importances, as in [Frères2016], also produced worse results than when using the permutation importance alone.

Given the fact that neither the random forests based gene ranking nor the extreme pseudo-samples based one outperformed the differential expression based one for all of the 12 data sets, one might wonder if using both a supervised learning based gene selection technique in conjunction with differential expression would not yield better results. However, using the supervised learning based gene selection method after the differential expression one (*i*.*e*. using only the genes with a significant differential expression adjusted p-value as input features of the random forests classifier or the EPS method) also produced worse results than when using the random forests gene ranking or the EPS gene ranking alone.

Using survival analysis as a way to validate gene lists coming from cancer data sets whose average survival differs greatly might spark questions, however there does not seem to be a link between the overall survival (OS) of these cancers and the performance of the proposed methods. Survival information constitutes a quantifiable and relatively easily available information for different data sets. However, using the presumed relationship between the expression values of a gene and the survival of a patient as a proxy for the role of said gene in the selected disease relies on a strong hypothesis whose validity might vary across data sets. Therefore, other gene ranking validation methods should be further explored to assess the performance of a random forests based gene ranking method and the EPS method in a wider range of RNA-Seq experiments.

In conclusion, we have shown that using the permutation importance internally computed by the random forests algorithm, when said algorithm is used to build a classifier based on gene expression values of a case-control RNA-Seq data set, allowed to obtain a ranking of genes; Variational Autoencoders could be used to generate pseudo-samples mimicking the properties of real samples, albeit with extreme localizations in latent space; Using the feature weights of said pseudo-samples allowed to obtain a ranking of genes. These rankings were compared with the results of a differential expression analysis, with all three gene rankings being evaluated through survival analysis on a validation cohort different from the cohort used to generate both rankings. The results have shown that the random forests based method and the extreme pseudo-samples outperformed the differential expression based method for 9 and 8 out of the 12 data sets analyzed, respectively. Although the genes selected by both methods are different, there is no significant difference in the number of highly correlated genes between both methods. Although the goal of this research is not to supersede differential expression analysis to select genes of interest in RNA-Seq studies, we have shown that differential expression analysis might miss out on important genes, and a supervised learning based gene selection method should be used alongside.

As the field of machine learning contains many different supervised classification and feature selection algorithms, it would be of interest to extend this work by testing the performance of other methods for gene selection in the context of case-control RNA-Seq data sets.

## 6 Author Contributions

SW: Conceived and designed the experiments; Performed the random forests analysis; Contributed to the writing of the manuscript. RS: Developed and performed the Extreme-Pseudo Samples analysis; Contributed to the writing of the manuscript

## 9 Acknowledgments

We thank Claire Josse, Pierre Geurts, Vincent Botta, Eimear Kenny, Gillian Belbin, Jose-Luis Ambite and Shunsuke Saito. This work was supported in part through the computational resources and staff expertise provided by Scientific Computing at the Icahn School of Medicine at Mount Sinai.

---

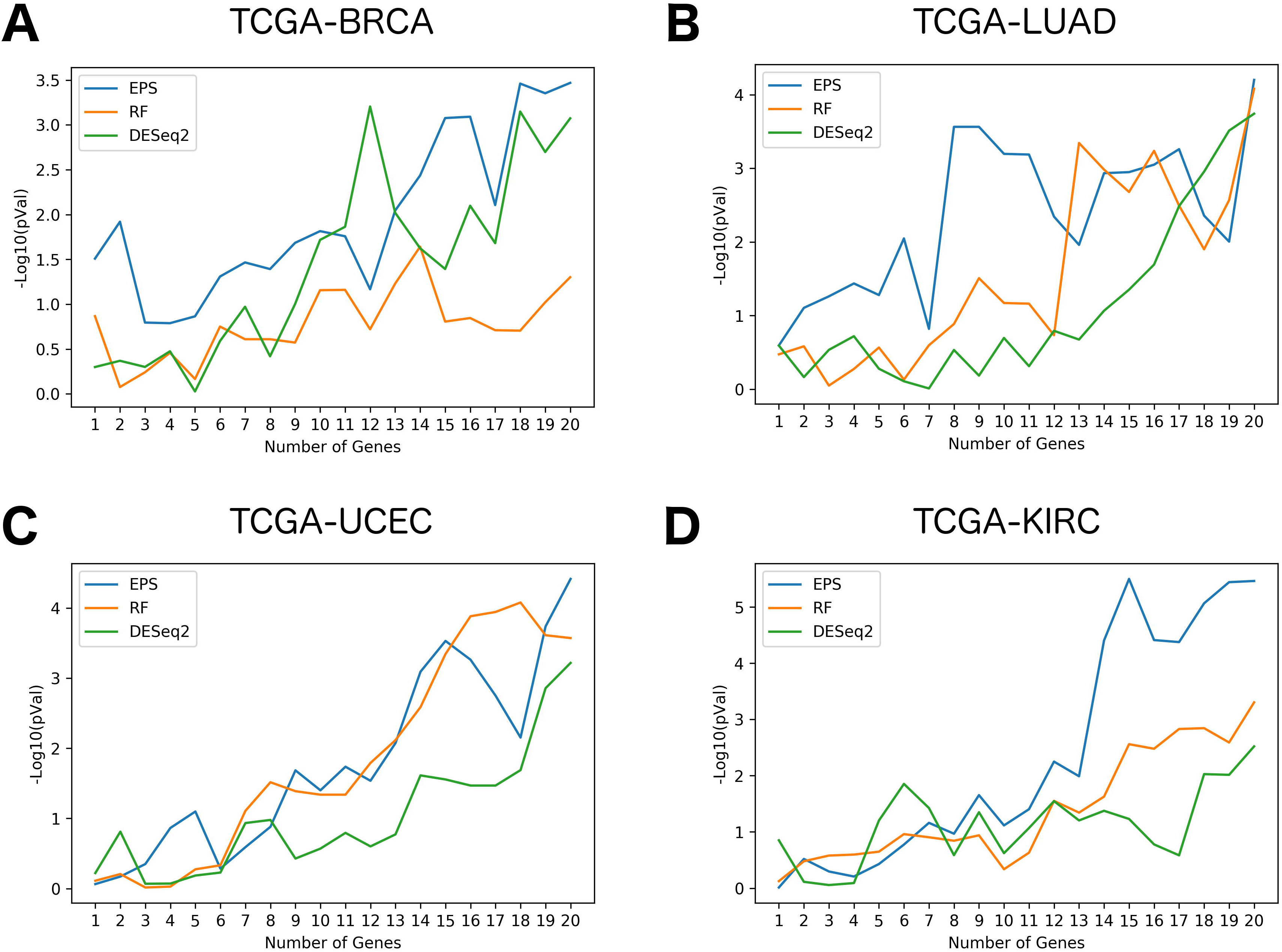

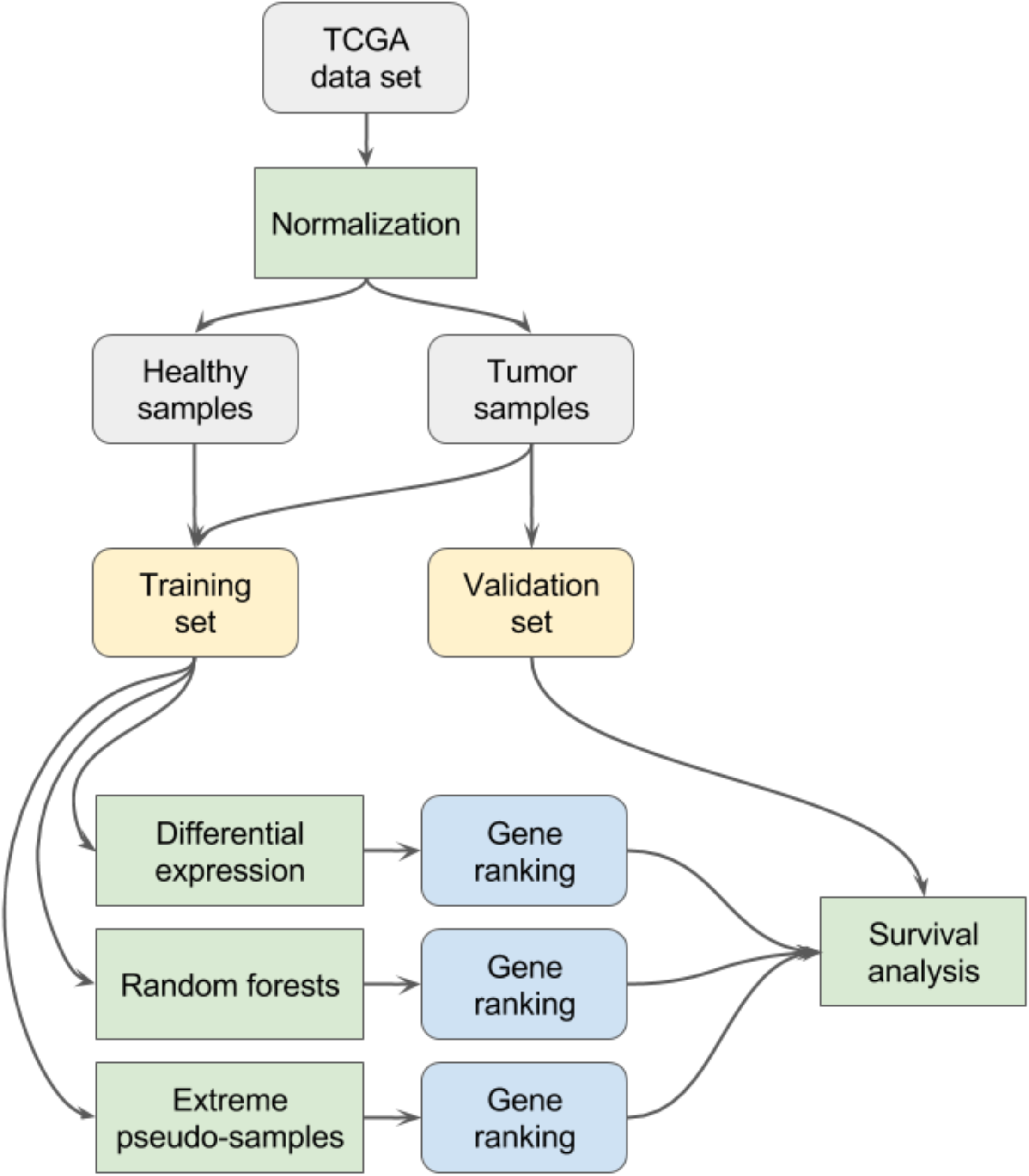

## References

1. Wang, Zhong, Mark Gerstein, and Michael Snyder. “RNA-Seq: a revolutionary tool for transcriptomics.” Nature reviews genetics 10, no. 1 (2009): 57.

2. Mortazavi, Ali, Brian A. Williams, Kenneth McCue, Lorian Schaeffer, and Barbara Wold. “Mapping and quantifying mammalian transcriptomes by RNA-Seq.” Nature methods 5, no. 7 (2008): 621.

3. Trapnell, Cole, Lior Pachter, and Steven L. Salzberg. “TopHat: discovering splice junctions with RNA-Seq.” Bioinformatics 25, no. 9 (2009): 1105–1111.

4. Garber, M., Grabherr, M. G., Guttman, M. & Trapnell, C. Garber, Manuel, Manfred G. Grabherr, Mitchell Guttman, and Cole Trapnell. “Computational methods for transcriptome annotation and quantification using RNA-seq.” Nature methods 8, no. 6 (2011): 469.

5. Love, Michael I., Wolfgang Huber, and Simon Anders. “Moderated estimation of fold change and dispersion for RNA-seq data with DESeq2.” Genome biology 15, no. 12 (2014): 550.

6. Wenric, Stephane, Sonia ElGuendi, Jean-Hubert Caberg, Warda Bezzaou, Corinne Fasquelle, Benoit Charloteaux, Latifa Karim et al. “Transcriptome-wide analysis of natural antisense transcripts shows their potential role in breast cancer.” Scientific reports 7, no. 1 (2017): 17452.

7. Huang, Huei-Chung, Yi Niu, and Li-Xuan Qin. “Differential Expression Analysis for RNA-Seq: An Overview of Statistical Methods and Computational Software: Supplementary Issue: Sequencing Platform Modeling and Analysis.” Cancer informatics 14 (2015): CIN-S21631.

8. Frères, Pierre, Stéphane Wenric, Meriem Boukerroucha, Corinne Fasquelle, Jérôme Thiry, Nicolas Bovy, Ingrid Struman et al. “Circulating microRNA-based screening tool for breast cancer.” Oncotarget 7, no. 5 (2016): 5416.

9. Yao, Dengju, Jing Yang, Xiaojuan Zhan, Xiaorong Zhan, and Zhiqiang Xie. “A novel random forests-based feature selection method for microarray expression data analysis.” International journal of data mining and bioinformatics 13, no. 1 (2015): 84–101.

10. Duro, Dennis C., Steven E. Franklin, and Monique G. Dubé. “Multi-scale object-based image analysis and feature selection of multi-sensor earth observation imagery using random forests.” International Journal of Remote Sensing 33, no. 14 (2012): 4502–4526.

11. Anaissi, Ali, Paul J. Kennedy, Madhu Goyal, and Daniel R. Catchpoole. “A balanced iterative random forest for gene selection from microarray data.” BMC bioinformatics 14, no. 1 (2013): 261.

12. Colaprico, Antonio, Tiago C. Silva, Catharina Olsen, Luciano Garofano, Claudia Cava, Davide Garolini, Thais S. Sabedot et al. “TCGAbiolinks: an R/Bioconductor package for integrative analysis of TCGA data.” Nucleic acids research 44, no. 8 (2015): e71–e71.

13. Weinstein, John N., Eric A. Collisson, Gordon B. Mills, Kenna R. Mills Shaw, Brad A. Ozenberger, Kyle Ellrott, Ilya Shmulevich, Chris Sander, Joshua M. Stuart, and Cancer Genome Atlas Research Network. “The cancer genome atlas pan-cancer analysis project.” Nature genetics 45, no. 10 (2013): 1113.

14. Wright, Marvin N., and Andreas Ziegler. “ranger: A fast implementation of random forests for high dimensional data in C++ and R.” arXiv preprint 1508.04409 (2015).

15. Vogelstein, Bert, Nickolas Papadopoulos, Victor E. Velculescu, Shibin Zhou, Luis A. Diaz, and Kenneth W. Kinzler. “Cancer genome landscapes.” science 339, no. 6127 (2013): 1546–1558.

16. Yates, Lucy R., and Peter J. Campbell. “Evolution of the cancer genome.” Nature Reviews Genetics 13, no. 11 (2012): 795.

17. Kanehisa, Minoru, and Susumu Goto. “KEGG: kyoto encyclopedia of genes and genomes.” Nucleic acids research 28, no. 1 (2000): 27–30.

18. Joshi-Tope, G., Marc Gillespie, Imre Vastrik, Peter D’Eustachio, Esther Schmidt, Bernard de Bono, Bijay Jassal et al. “Reactome: a knowledgebase of biological pathways.” Nucleic acids research 33, no. Suppl_1 (2005): D428–D432.

19. Phillips, Patrick C. “Epistasis—the essential role of gene interactions in the structure and evolution of genetic systems.” Nature Reviews Genetics 9, no. 11 (2008): 855.

20. Vidal, Marc, Michael E. Cusick, and Albert-László Barabási. “Interactome networks and human disease.” Cell 144, no. 6 (2011): 986–998.

21. Kingma, Diederik P., and Max Welling. “Auto-encoding variational bayes.” arXiv preprint 1312.6114 (2013).

22. Tan, Jie, Matthew Ung, Chao Cheng, and Casey S. Greene. “Unsupervised feature construction and knowledge extraction from genome-wide assays of breast cancer with denoising autoencoders.” In Pacific Symposium on Biocomputing Co-Chairs, pp. 132–143. 2014.

23. Danaee, Padideh, Reza Ghaeini, and David A. Hendrix. “A deep learning approach for cancer detection and relevant gene identification.” In PACIFIC SYMPOSIUM ON BIOCOMPUTING 2017, pp. 219–229. 2017.

