## Supplementary figures and images for "Using supervised learning methods for gene selection in RNA-Seq case-control studies"

### Supplementary Materials

TCGA-HNSC

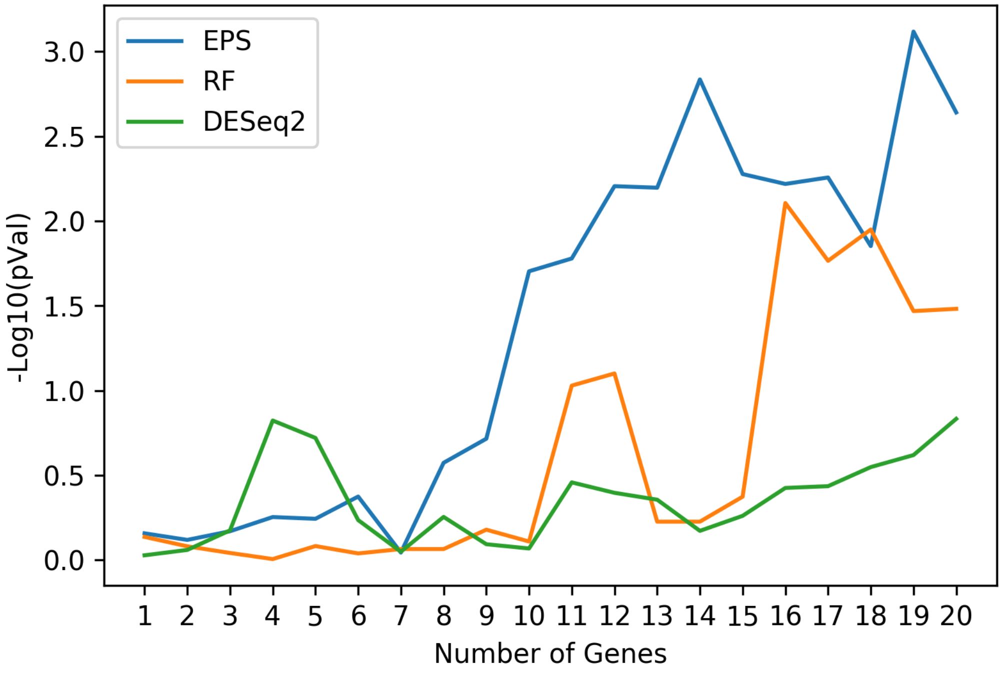


TCGA-THCA

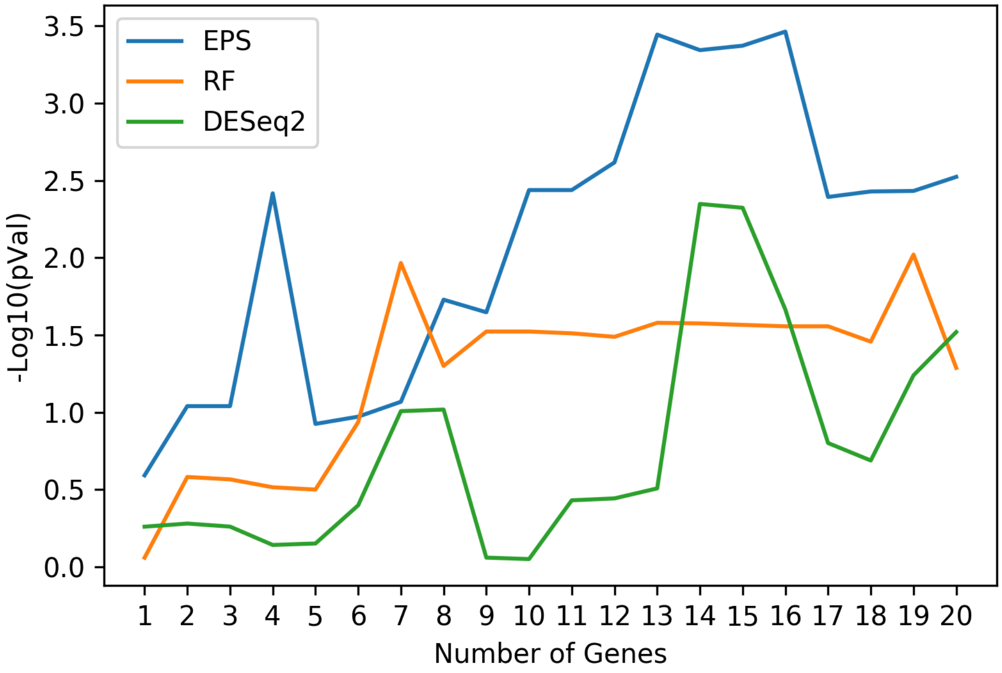

TCGA-LUSC

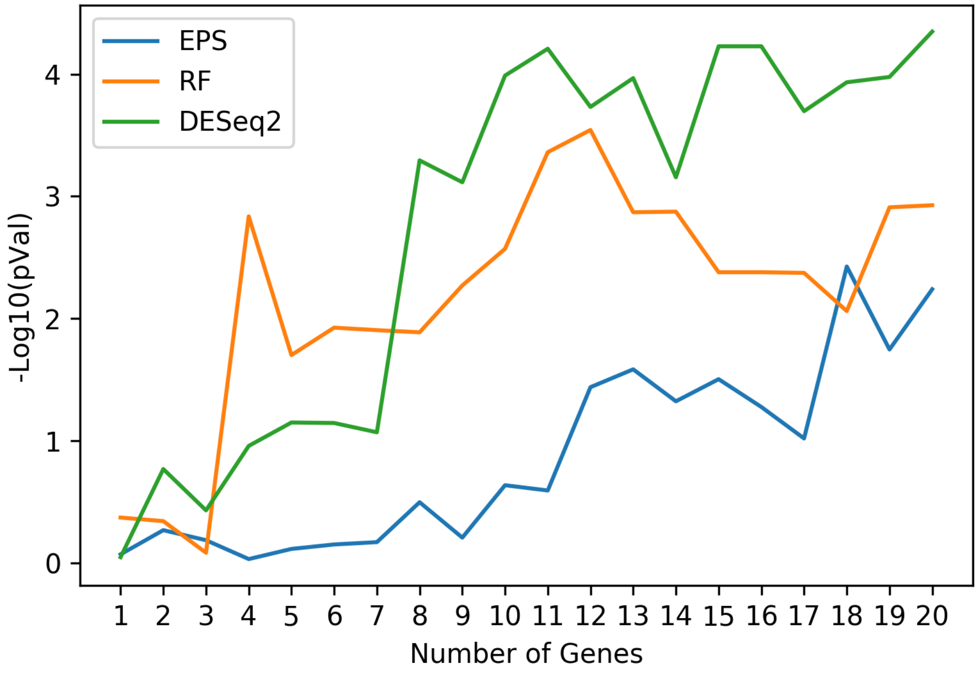


TCGA-PRAD

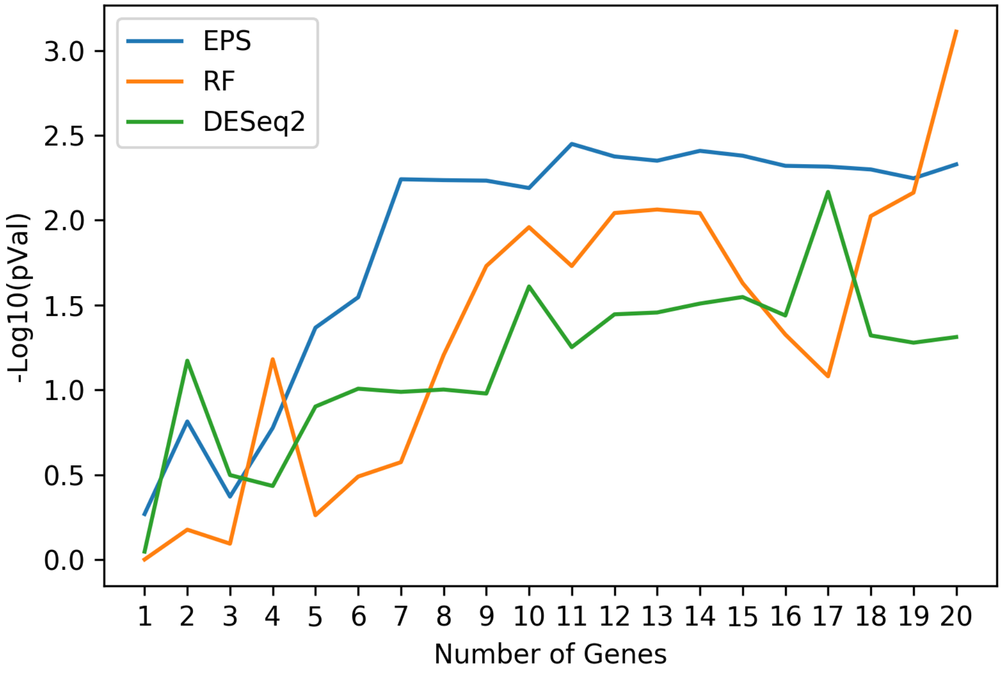

TCGA-COAD

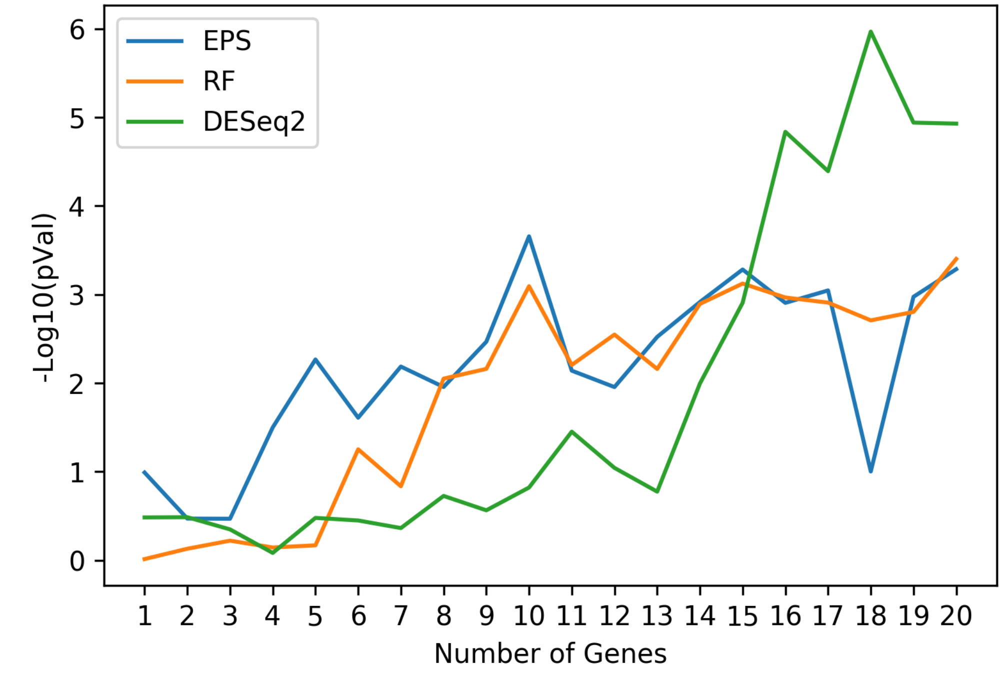


TCGA-STAD

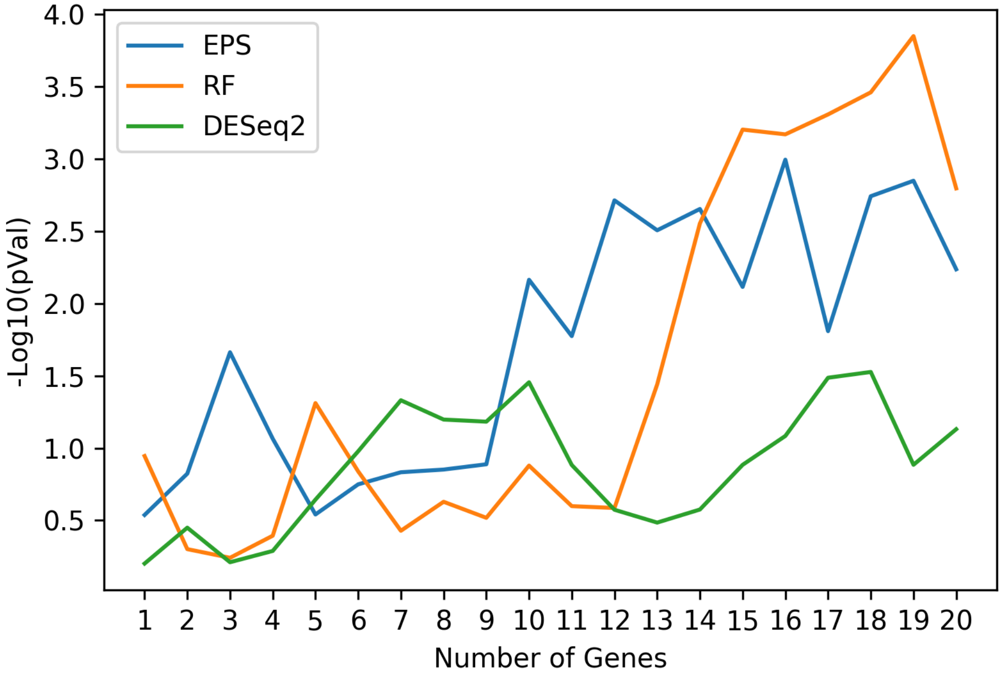

TCGA-LIHC

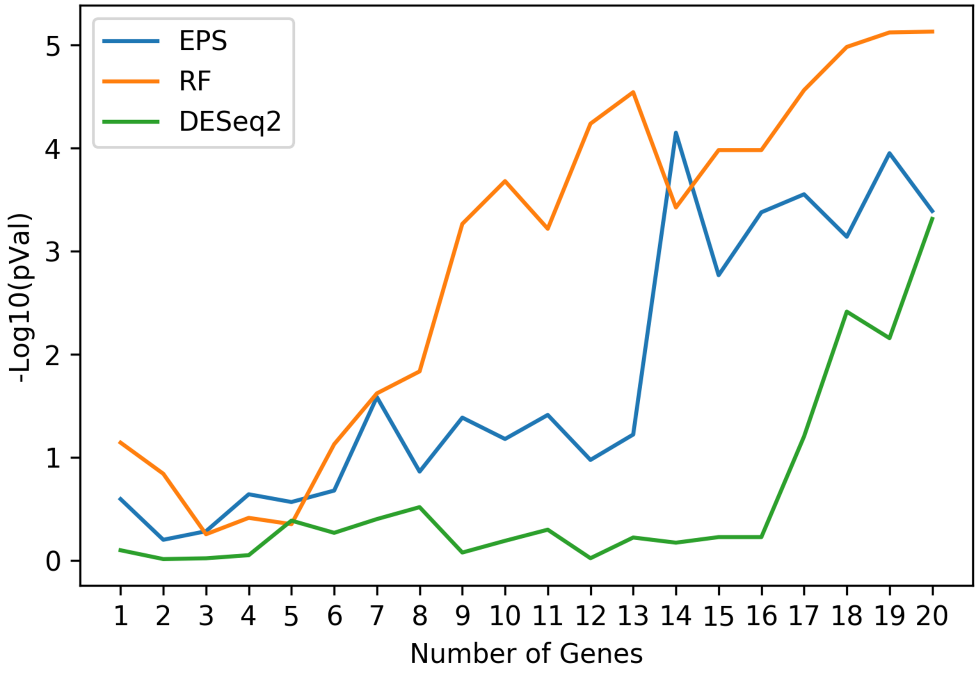


TCGA-KIRP

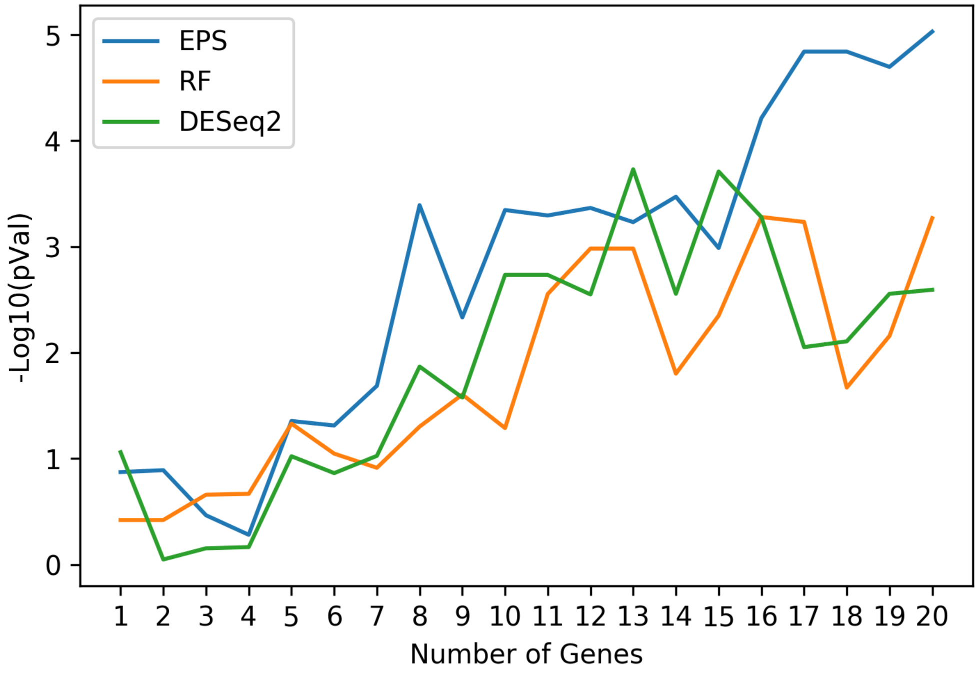
